# Classification and analysis of retinal interneurons by computational structure under natural scenes

**DOI:** 10.1101/2024.03.18.585364

**Authors:** Dongsoo Lee, Juyoung Kim, Stephen A. Baccus

## Abstract

Inhibitory neurons are diverse across the brain, but for the visual system we lack the ability to functionally classify these neurons under complex natural stimuli. Here we take the approach of classifying retinal amacrine cell responses to natural scenes using optical recording and an interpretable neural network model. We fit mouse amacrine cell responses to a two-layer convolutional neural network model of a class shown previously to accurately capture salamander ganglion cell responses to natural scenes. Using an approach from interpretable machine learning, we determined for each stimulus the model interneurons that generated each amacrine response, analogous to the set of bipolar cells that target the amacrine population. From this analysis we clustered amacrine cells not by their natural scene responses, but by the model presynaptic neurons that constructed those responses, conservatively finding approximately seven groups by this approach. By analyzing the set of model presynaptic input neurons for each amacrine cluster, we find that distributed rather than dedicated inputs generate natural scene responses for different amacrine cell types. Additional analyses revealed distinct transient and sustained modes exhibited by the network during the response to simple flashes. These results give insight into the computational structure of how the diverse amacrine cell population responds to natural scenes, and generate multiple quantitative hypotheses for how synaptic inputs generate those responses.

## Introduction

Visual computation in the mammalian retina is implemented by a large set of interneurons, the most diverse being inhibitory amacrine cells (Masland, 2012). Although there has been substantial recent progress in classifying amacrine cell types based on their molecular expression (Yan et al., 2020), we lack approaches to functionally classify diverse sets of neurons under natural stimuli. Several factors make this problem challenging, including distributed anatomical connectivity (Helmstaedter et al., 2013) and the high level of nonlinearity in the retina (Gollisch and Meister, 2010), which makes visual sensitivity stimulus-dependent.

Previous methods to functionally classify cells include classification directly from responses (Baden et al., 2016), which is primarily useful when cells receive identical stimuli as in the case of a uniform field or moving bar. Complex spatiotemporal stimuli, however, pose a problem because the stimulus is different for each cell by virtue of their different position, pointing to the use of a computational model with which to capture and compare response properties. White noise analysis has been useful to classify neurons by their simple spatiotemporal sensitivity and threshold properties (Segev et al., 2006), but only does so under specific stimulus statistics.

For natural scenes, previously we have successfully captured the responses of salamander retinal ganglion cells with a three-layer convolutional neural network model (McIntosh et al., 2016) (Maheswaranathan et al., 2023). In addition to accurately capturing the natural scene response, we find these models have two critically important properties 1) Generalization, meaning that fit only to natural scenes, the models successfully replicate a wide range of artificial phenomena. 2) Interpretability, meaning that the activity of internal units is highly correlated with measured interneuron recordings that the model was never fit to (Maheswaranathan et al., 2023).This demonstrates that CNNs not only accurately capture sensory responses to natural scenes, but also can yield information about the circuit’s internal structure and function.

An additional useful aspect of these models is that they can be analyzed to determine the set of model interneurons that generated any particular response (Tanaka et al., 2019) (Maheswaranathan et al., 2023). This analysis, termed an attribution analysis, derives from approaches of interpretable machine learning (Sundararajan et al., 2017), where the goal is to gain insight into how the properties of the model generated particular outputs. Because the interneurons within the model bear functional similarity to real interneurons, determining which model interneurons contributed to model output automatically generates hypotheses as to which real interneurons caused a particular response.

Here we classify hundreds of mouse amacrine cells recorded simultaneously using calcium imaging from cells expressing GCaMP6f responding to flashes and natural scenes. We fit two-layer convolutional neural network models to amacrine cell responses. Using the method of Integrated Gradients, we calculate the contribution of every Layer 1 neuron to every amacrine cell for every stimulus, representing the set of presynaptic cells that drove each separate response. We classified amacrine cells by these contributions identifying at least seven distinct classes. An analysis of this set of contributions indicated that rather than dedicated parallel pathways, input to amacrine cells was distributed, with multiple presynaptic cell types influencing multiple different amacrine types in all cases. Given that the interneurons of this class of models has been shown to have high correlations with real interneurons, it is expected that this approach will yield not only a method of classification, but one that automatically yields testable quantitative hypotheses as to the presynaptic circuits that generate the diversity of amacrine cell responses under natural scenes.

## Results

We recorded optical responses from mouse amacrine cells using a custom resonant scanning two-photon microscope by expressing GCaMP6f under control of the molecular marker, Ptf1a-Cre (Fig. 1). Stimuli included uniform field flashes, spatiotemporal white noise, and natural scenes consisting of natural movies jittered with the statistics of eye movements. Hundreds of regions of interest (ROIs) were identified by grouping nearby pixels that shared a highly correlated time course. For natural scenes, ROIs were accepted that had a sufficiently high correlation between repeats of the stimulus. To identify different types of amacrine cells by a standard method, we classified cells by their flash responses, finding approximately 6 classes (Fig. 2). These included On cells, Off cells and On-Off cells having different time courses.

**Figure 1.**
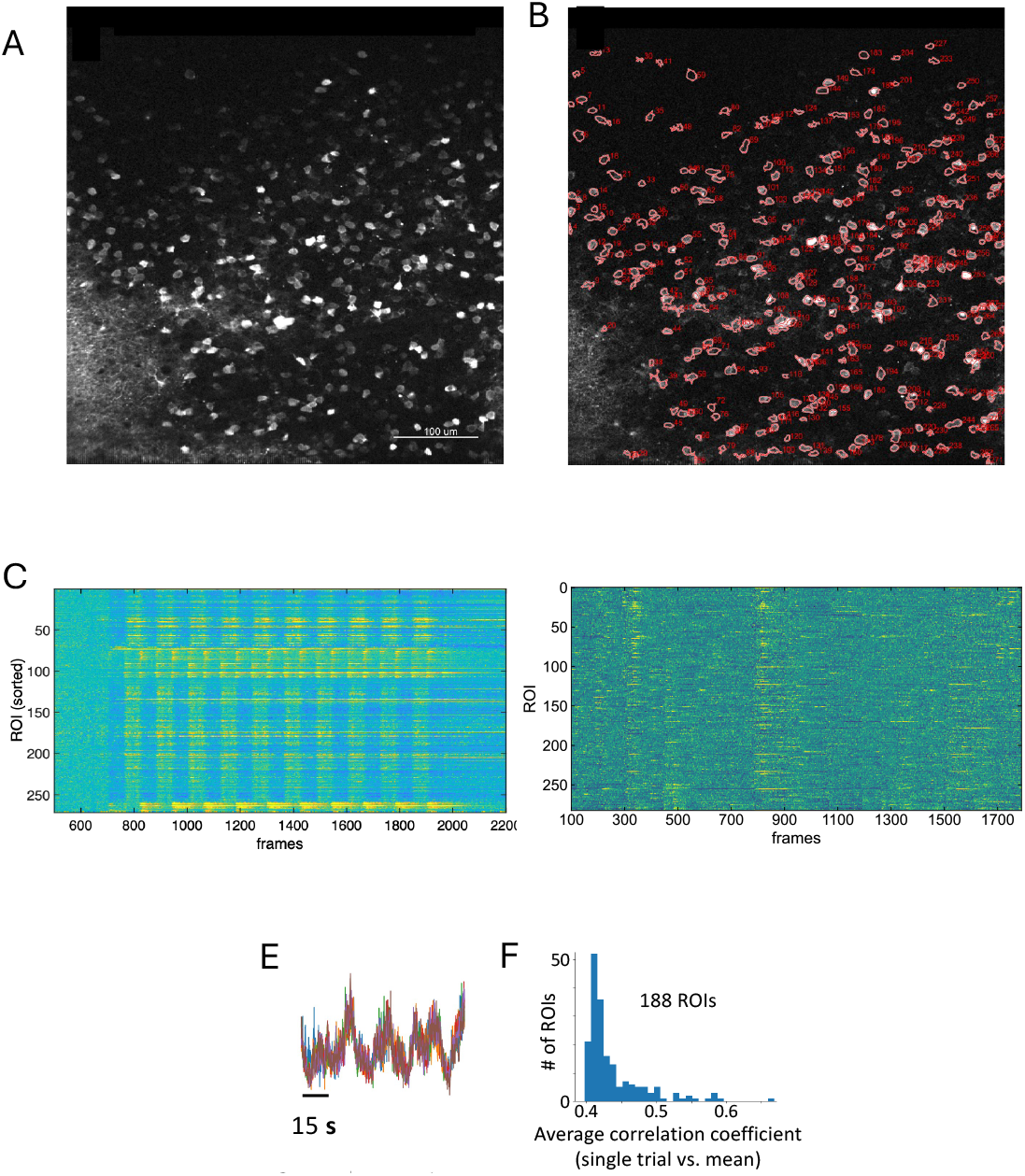
Amacrine cell population responses to flashes and natural scenes. Calcium responses were recorded from all amacrine cell types using a Ptf1a-CRE and a GCaMP6f reporter. A. Wide field image. B. Identified regions of interest (ROIs). C. Flash responses recorded in a different preparation. Each row is a different cell, grouped by clustering flash responses using k-means. D. Example set of responses to a set of natural scenes, with no cell grouping. E. Repeated presentation of a natural movie for a single ROI. F. Histogram of average correlation coefficient between a single trial and the mean response for a population of cells.

**Figure 2.**
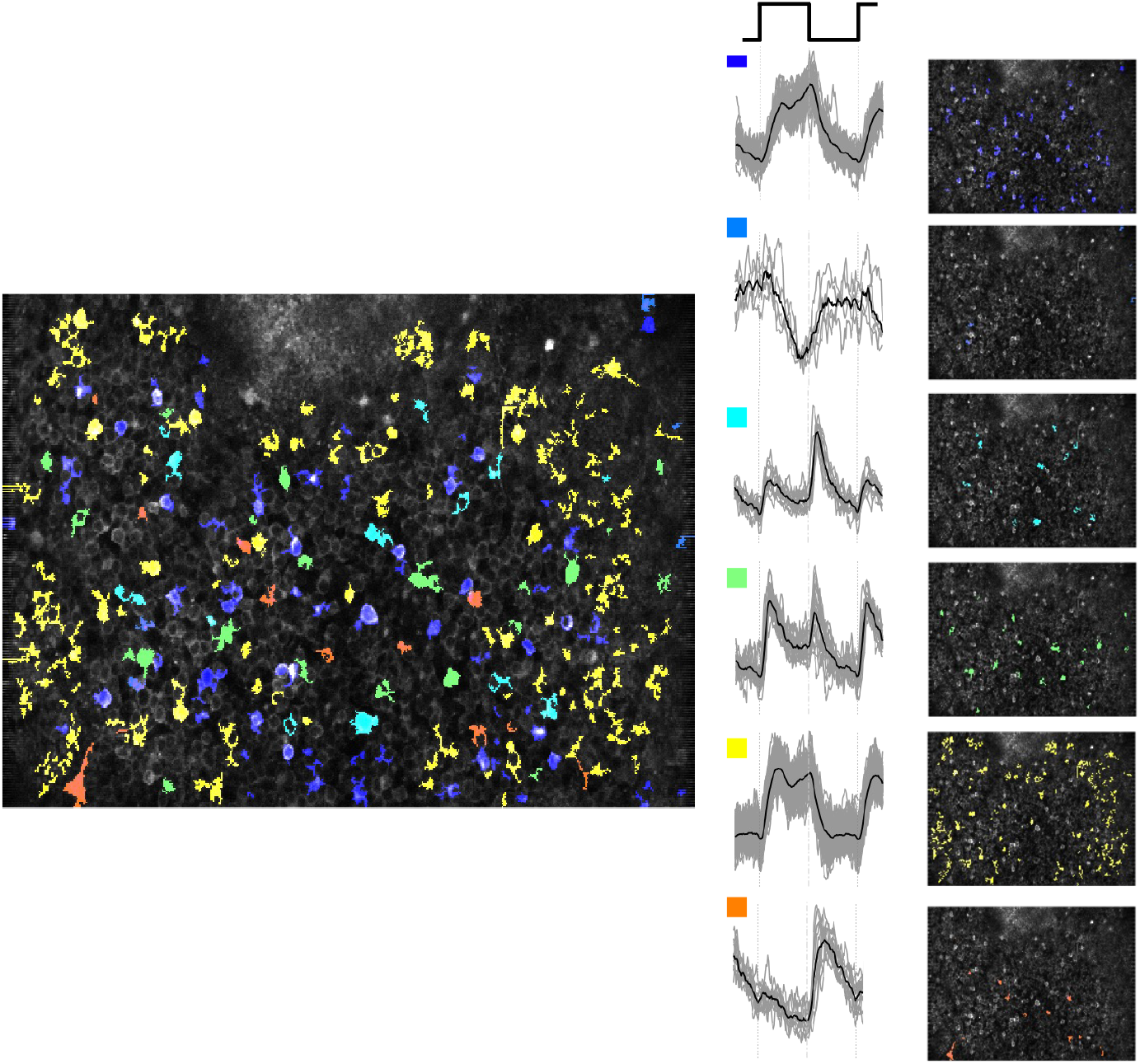
Clusters of amacrine cell flash responses. A. Amacrine cells clustered by their flash responses (3 s on, 3 off). B. Superimposed responses from many cell (grey traces) and the mean of the cluster (black trace). C. Spatial locations of cells from each cluster.

We then fit a two-layer CNN to the natural scene responses of a population of amacrine cells (Fig. 3a). Two layers were chosen because of the previous success of three layer models of ganglion cells (McIntosh et al., 2016). Layer 1 was convolutional representing a mosaic of interneurons, and had six model interneurons types that we would expect to correspond to bipolar cells. The second layer was a fully connected layer that received spatially weighted input from each of the Layer 1 cell types.

**Figure 3.**
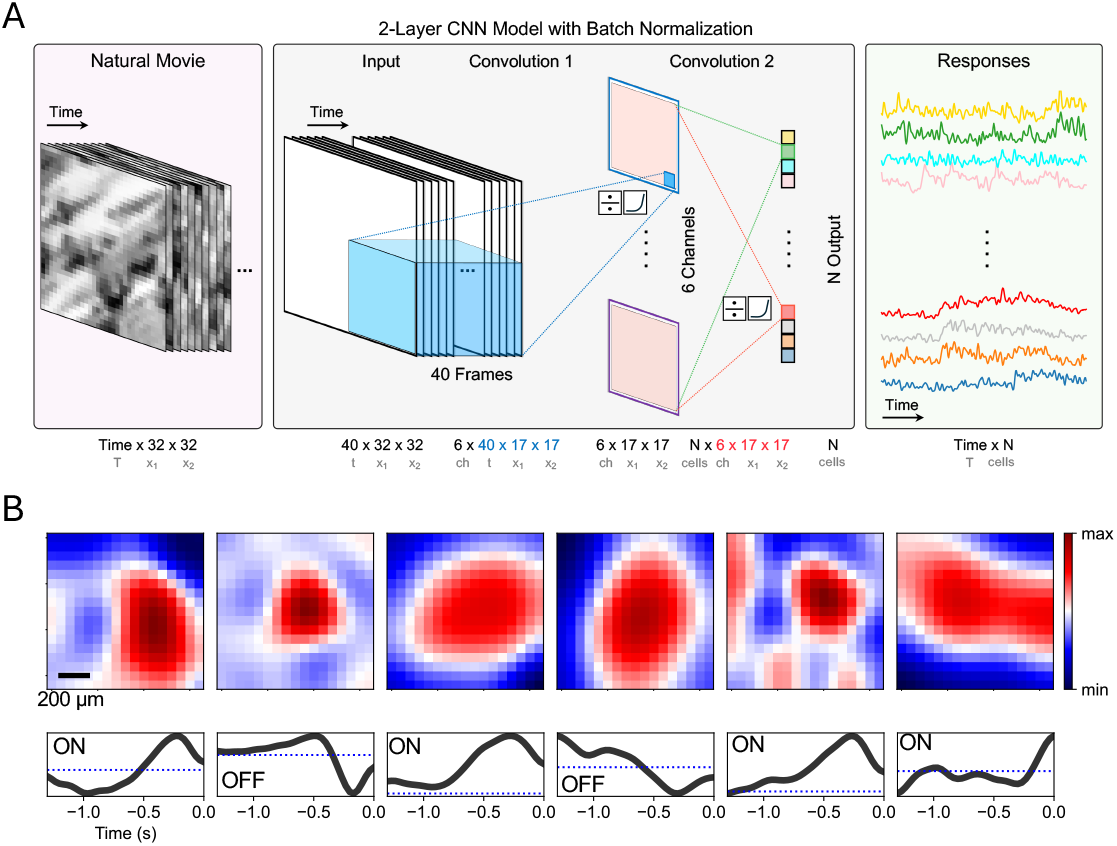
Convolutional neural network model of amacrine cell responses. The model is two layers, with 6 cell types in the first layer, analogous to bipolar cells, and a second layer that combines weighted input from the six Layer 1 cell types. A. Diagram of two-layer CNN model. B. Spatial and temporal components of the spatiotemporal receptive field, shown as the separable (Rank-1) decomposition of the receptive field, although the full non-separable receptive field is fit by the model. The sign of the receptive field (On vs. Off) is represented and labelled in the time course.

The receptive fields of Layer 1 cells were of different sizes and time courses, including On and Off cells, and monophasic (sustained) and biphasic (transient) cells as expected (Fig. 3b). Although more bipolar cell types exist than six, the goal here was not to replicate the full set of known individual bipolar cells types, but rather to create a minimal model that captured the amacrine cell data. Thus, it is likely that some Layer 1 cell types approximated the combined input created by more than one bipolar cell. In particular, signs of the Layer 1 cells were not constrained to be of one sign (known as Dale’s law), and thus could be both positive or negative. If this constraint were added, such an arrangement would be produced by two different excitatory bipolar cells, one On, and one Off.

We sought to classify amacrine cells by the structure of the model neural pathways that caused their response, analogous to classifying them according to the bipolar cells that drove their responses. Previously, we defined the interneuron contribution (INC) as the net excitation or inhibition that each model interneuron makes to each cell of the model’s output, for each particular stimulus (Maheswaranathan et al., 2023). This contribution is a function of two properties, the amount that the interneuron is driven by the stimulus, and the effect of that interneuron on the model’s output given the particular stimulus (Fig. 4a). This quantity is stimulus-dependent because the circuit is nonlinear, meaning that in the general case an interneuron may respond with low or high sensitivity to the current stimulus (although in the present case Layer 1 neurons are linear), and the pathway between the interneuron and the model output may also be in a state of low or high sensitivity because of intervening nonlinearities.

**Figure 4.**
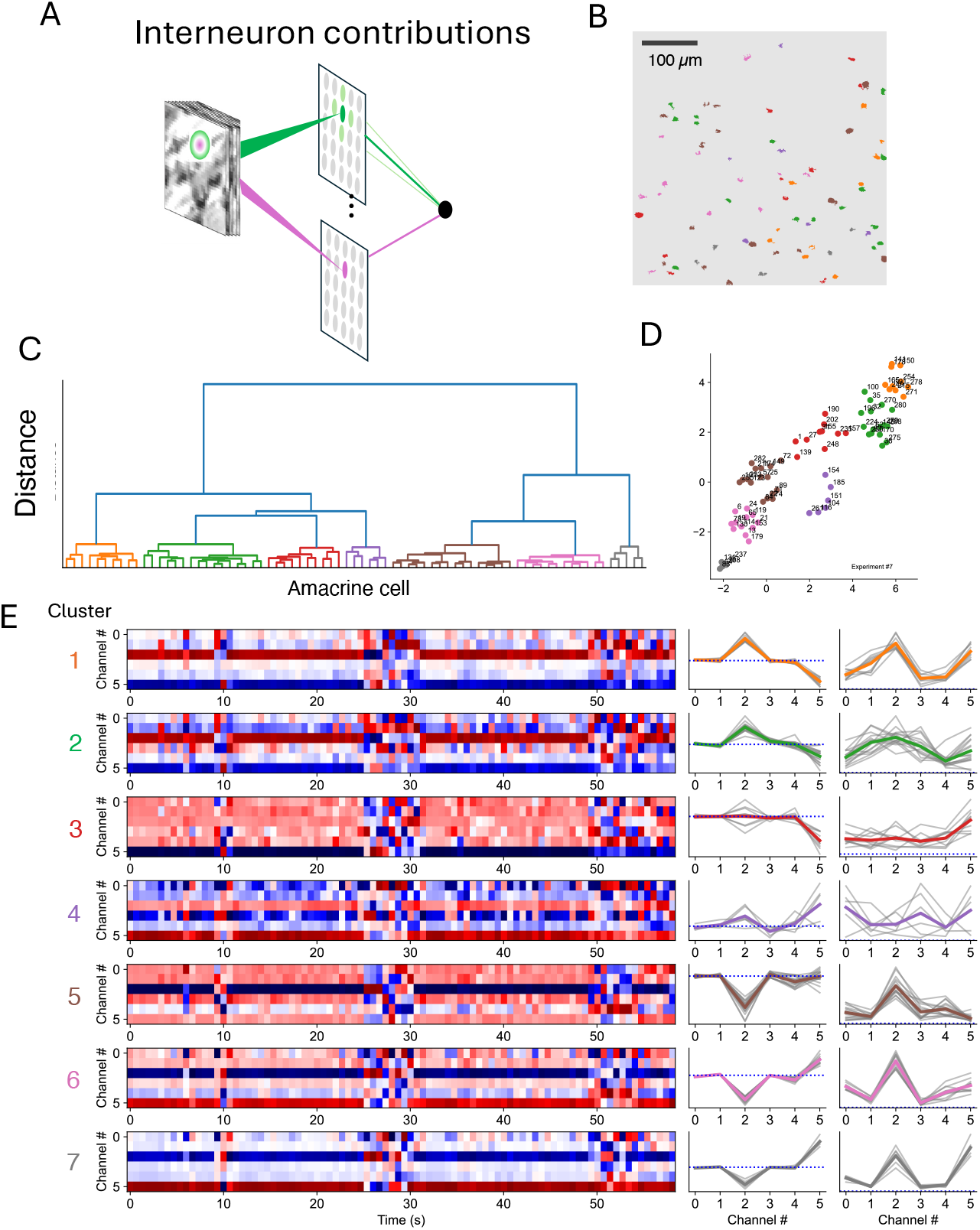
Analysis of interneuron contributions. A. Diagram of concept of interneuron contributions. B. Spatial locations of ROIs. C. Hierarchical clustering diagram of Layer 1 INCs that caused amacrine cell responses to natural scenes. D. tSNE plot showing individual ROIs and assigned clusters. E. INCs for each amacrine cell cluster in response to a 60 s segment of natural scenes. Left. Each row within each amacrine clusters response is a different Layer 1 cell type (Channels 0 – 5). Red indicates excitation, blue indicates inhibition. Middle. Average INC for each Layer 1 cell type for each cluster. Right. Standard deviation of INC for each cluster, showing the magnitude of the contribution of each Layer 1 cell to each amacrine cluster.

To compute the INCs for each stimulus, we used the method of Integrated Gradients (Sundararajan et al., 2017) (Tanaka et al., 2019) which computes the average contribution of the interneuron along a straight path from a zero stimulus (a grey screen) to the current stimulus, *s*. This quantity represents the average contribution of the interneuron as the stimulus changes from zero to *s*. A straight path is chosen in order for the method to be sensitive to any interneuron that causes a change in the response, and for the method to be invariant to the particular implementation of the intervening network (Sundararajan et al., 2017).

We computed the contribution of each Layer 1 interneuron to each amacrine cell at each time point. We then clustered amacrine cells by their average INCs from Layer 1, representing the relative contribution of each of the six Layer 1 cell types under natural scenes. We identified 5 – 7 amacrine cell clusters that had differing computational structure as measured by the contributions from Layer 1 cell types (Fig. 4, Supp. Fig 1). For each amacrine cluster, these INCs differed in their mean input (excitatory or inhibitory) and standard deviation. The standard deviation of the INC across a natural scene segment represented the magnitude of the contribution from each Layer 1 interneuron to each amacrine cluster, forming a matrix of contributions (Fig. 5a). This contribution matrix is to be distinguished from a synaptic weight matrix, which would simply account for the connection weight from Layer 1 cell to amacrine cell. The contribution matrix accounts also for how much each Layer 1 interneuron was activated by the particular set of stimuli and is thus stimulus dependent.

**Figure 5.**
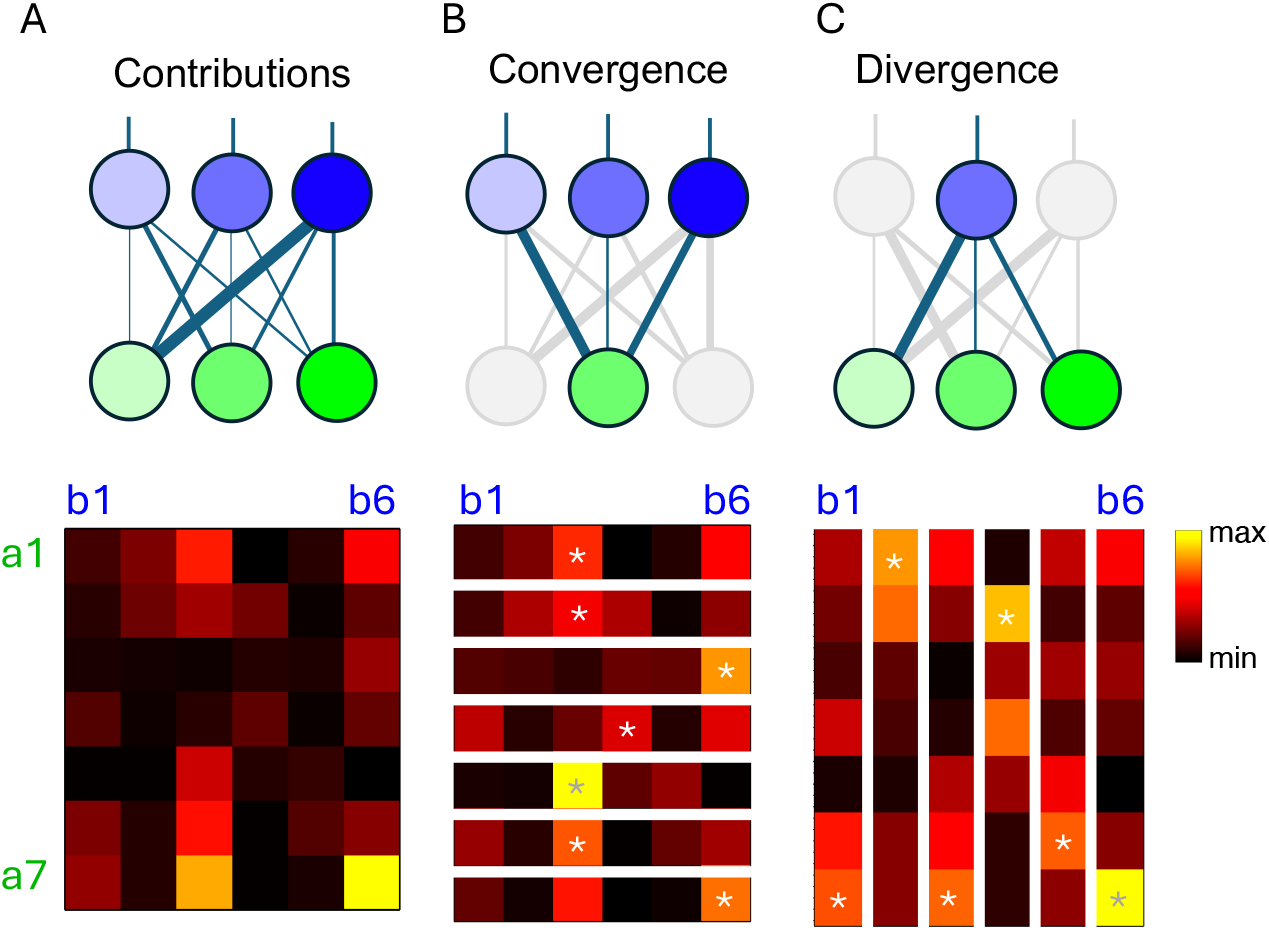
Structure of Interneuron Contributions to amacrine responses under natural scenes. A. Top. Diagram indicating the unnormalized contributions of Layer 1 interneurons (blue) to drive amacrine cells (green). Bottom. Matrix of average INC contributions for each cluster, Layer 1 cell types are labelled b1 – b6, amacrine cell clusters ar a1 – a7. B. Top. Diagram indicating analysis of how presynaptic inputs converge to postsynaptic inputs. Bottom. Normalized matrix by rows showing convergence of Layer 1 neurons for each amacrine cluster. Each row has the same total weight, which indicates the presynaptic cells that drove each postsynaptic cell the most (asterisk). C. Top. Diagram indicating divergence of single presynaptic cell types to the set of postsynaptic cell types. Bottom. Normalized matrix by columns showing divergence of output to each amacrine cluster for each Layer 1 neuron type. Asterisk indicates the amacrine cell driven the most, on average, during natural scenes by each Layer 1 cell type. For only one cell type pair do the greatest input and the greatest effect coincide (bottom right, b6 – a7), which is the minimum possible given that the maximum overall contribution must be the maximum of both row and column.

We analyzed this matrix of contributions first by normalizing the rows, which indicated for each amacrine cell cluster the relative weighting of each Layer 1 cell that contributed to that amacrine cluster. From this representation, we could determine which Layer 1 cell had the greatest contribution to each amacrine cell type (Fig. 5b). An alternative analysis normalized the columns of the matrix, indicating for each Layer 1 cell the relative effect on each amacrine cell type. This revealed for each Layer 1 cell which amacrine cell type was affected the most (Fig. 5c). The concept of parallel pathways or a labelled line can be quantitatively defined by cases where an input cell type and output cell type choose each other to have the strongest output and input, respectively. In other words, where the row and column of the contribution matrix have the same maximum. The extreme case of parallel pathways is represented by a diagonal matrix, where each presynaptic cell type is dedicated to only a single postsynaptic cell type, and vice versa. Examining the contribution matrices normalized by rows and columns, this coincidence of greatest effect and greatest input occurred in only one cell type pair in the contribution matrix, which is the minimum possible given that the overall maximum must be both the maximum of its row and column. Therefore, we conclude that overall, under natural scenes, the contributions from Layer 1 cell types to amacrine cell types conformed to a distributed, not a parallel organization.

We then compared the flash responses of the model fit to natural scenes to the flash responses recorded directly from the amacrine population. Flash responses of different amacrine cell types included On, Off, and On-Off responses of different time courses, with cells in individual clusters having similar responses (Fig. 6a). The CNN model was never directly fit to the recorded flash responses. We observed a close match between model and recorded flash responses. Although this match was not one to one, both flash response clustering and natural scene clustering could produce slightly different results with different decision criteria. Given that different stimuli are expected to drive different sets of amacrine inputs, it was not expected that the match be exact. Examining the interneuron contributions for the flash response for each amacrine cell type cluster of the model, we found that different Layer 1 interneurons changed their contribution dynamically during the flash response, and that different clusters had a different set of interneurons that contributed, as was the case for natural scenes.

**Figure 6.**
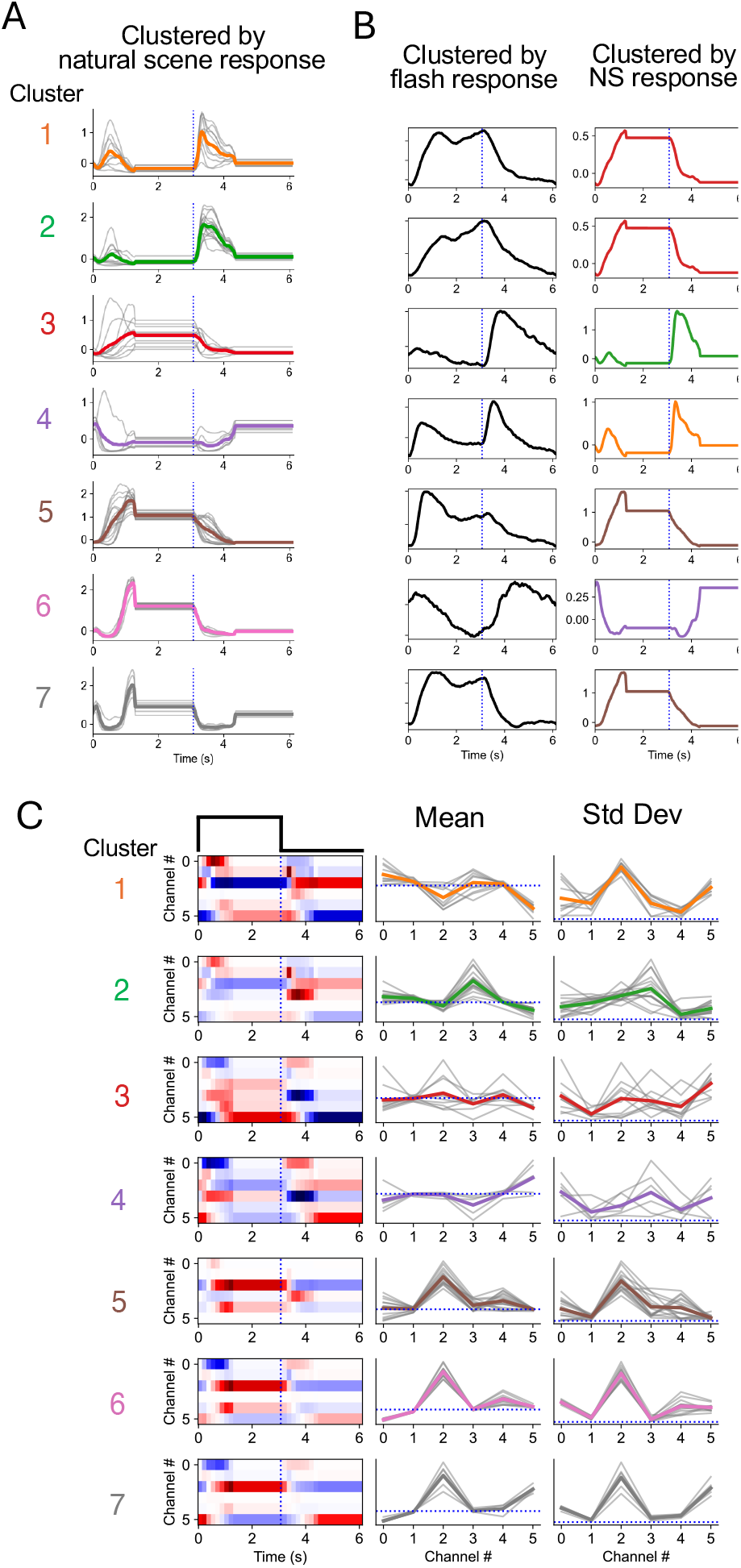
Interneuron contributions during flash response. A. Responses to a flash presented to the model for each model interneuron amacrine cluster. Clusters were found from natural scene responses. B. Comparison of recorded flash responses and model flash responses. Left. Average flash responses computed directly from flash responses. Right. Best match to amacrine clusters computed from natural scene responses from (A). C. INCs during flash responses for each amacrine cluster computed from natural scenes. Left. For each cluster, each row is the INC from each Layer 1 cell. Red indicates excitation, blue indicates inhibition. Middle, Colored traces indicate average INC for each cluster. Grey traces indicate average INC for individual amacrine cells in each cluster. Right, Standard deviation of INCs for each cluster. Colored traces indicate the average s.d. for Layer 1 INCs for each cluster, and grey traces indicate the s.d. of INCs computed across the flash response for each amacrine cell. The s.d. indicates the amplitude of the effect of each Layer 1 cell type to each amacrine cell.

Given that different interneurons contributed at different times, we then analyzed the overall state of the entire network at different times relative to the flash response (Fig. 7). Taking the set of INCs from all Layer 1 cells to all amacrine cell types, for the model with 7 amacrine types and 6 Layer 1 cells, at each point in time there were 42 INCs when averaged over all cells of a given type. Treating these as a vector, we clustered these to identify different states or modes of the network at each point in time. For the flash response, a hierarchical clustering analysis showed four different modes, although the exact number was uncertain (Fig. 7C). When we examined the timing of these modes, we observed a transient and sustained mode that occurred when the flash turned on, and a transient and sustained mode that occurred when the flash turned off (Fig. 7D). Overall, these modes showed different states of how the network constructed the flash response, and indicated the interneurons primarily responsible for the transient and sustatined states of the response.

**Figure 7.**
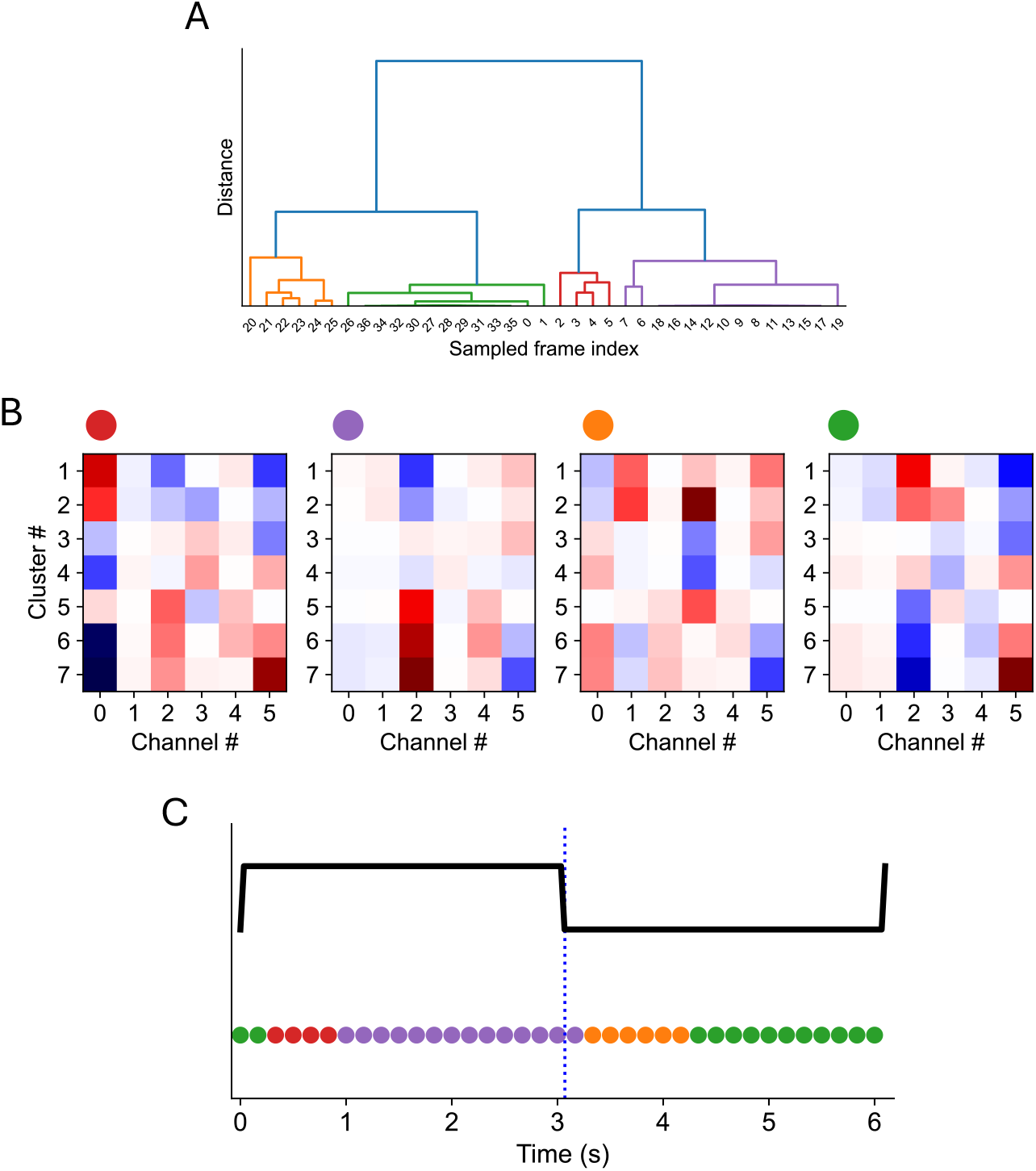
Modes of the network during a simple flash. At each time point, all INCs across all amacrine cell clusters were represented as a single vector and clustered to determine states or modes of activity across the network. A. Cluster analysis of modes of INCs during a flash response. Four clusters were chosen. B. The INC matrix for the four clusters, indicated the contribution of each Layer 1 cell type to each amacrine cell type. Red indicates excitation, blue indicates inhibition. C. Timing of four different modes compared to the flash responses. Colored symbols correspond to symbols in (B), indicating that the time course that the different modes occurred. Two of the modes occurred transiently after the stimulus changed, and two were more sustained, occurring with a delay.

## Discussion

Here we have taken a new approach to classify a diverse population of neurons using the computational structure of a network model. To classify a set of sensory neurons based on their functional properties, one must first choose a particular stimulus set. A network model that accurately captures responses to a wide range of stimuli is useful in that it allows a broad set of stimuli to be considered. Such a model allows a classification based not only on the model responses, but also on the internal states of the network that drove those responses. We have used a network model to classify a large set of amacrine cells based on the model interneurons that caused the responses to specific stimuli. Nonlinearities in the circuit make it especially important to consider the specific stimulus, as the set of bipolar cells that drive the response will likely differ for different stimuli.

In the salamander, model interneurons in a CNN model fit to ganglion cell responses to natural scenes have been shown to be highly correlated with real interneurons recorded separately (Maheswaranathan et al., 2023). This raises the possibility that classification by computational neural pathways can directly generate circuit hypotheses as to the set of bipolar cells that provide the effective input to different amacrine types.

The current model aimed to create a model with a minimum number of cell types to create amacrine responses under natural scenes. To make the more closely match retinal circuitry, one natural improvement is to constrain inputs to amacrine cell to be positive, as is the case with bipolar cells. This constraint would likely increase the number of required Layer 1 cells, causing positive and negative input to an amacrine cell from a single Layer 1 cell to be created by a combination of On and Off cells.

Known examples of specific connections to amacrine or ganglion cells involve dedicated inputs, such as the rod bipolar cell to AII amacrine cell (Vaney et al., 1991) or midget bipolar cells to midget ganglion cells in the primate (Kolb and Dekorver, 1991). Monostratified amacrine cells may receive similar dedicated input from a small number of bipolar cell types (Masland, 2012). However, other amacrine cells span multiple layers in the inner plexiform layer, and receive input from multiple bipolar cell types. In this case, one expects high divergence and convergence from the set of bipolar cells to the set of amacrine cells, as supported by connectomic studies (Helmstaedter et al., 2013). In our analysis, the contributions from the first layer to the different classes of amacrine cells conformed to this distributed organization, rather than dedicated parallel pathways. This distributed functional connectivity makes it particularly important to have quantitative hypotheses and measurements of synaptic effects. Current connectomic measurements are usually all or none, elevating the importance for causal experimental approaches perturbing bipolar cells to quantitatively measure the resulting functional effects on different amacrine for different stimuli. The interneuron contributions described here that can computed for any stimulus form the basis of such quantitative hypotheses, providing a guide for future experiments to reveal the circuit basis for the diverse set of inhibitory cell types in the retina.

## Methods

### Animals

All experimental protocols were approved by the Stanford University Institutional Animal Care and Use Committee and followed the guidelines of the National Institutes of Health. The offspring of heterozygous Ptf1a-Cre mice (B6.129S6(Cg)-*Ptf1a*^*tm2(cre/ESR1)Cvw*^/J, strain #036577, Jackson Laboratory) crossed with Ai148 mice (B6.Cg-*Igs7*^*tm148*.*1(tetO-GCaMP6f,CAG-tTA2)Hze*^/J, strain #030328, Jackson Laboratory) were used for optical imaging experiments. All mice were maintained on a C57BL/6J background (strain #000664, Jackson Laboratory). Mice were housed in groups and given food and water *ad libitum*. No statistical methods were used to pre-determine sample sizes.

### Tissue preparation

Mice of either sex were dark adapted for more than 2 hours prior to experiments. All tissue preparation was performed under dim red light. After euthanasia, both eyes were enucleated and hemisected along the cornea with scissors (Fine Science Tools). One eyecup was dissected at room temperature in Ames’ medium (A1420, Sigma-Aldrich) buffered with sodium bicarbonate (Fisher Scientific), equilibrated with 95% O_2_ and 5% CO_2_ (Praxair) at 100 mL/min, while the other eyecup was stored in the same solution. Dissection was performed with infrared LED illumination (>850 nm) under a stereomicroscope (MZ95, Leica Microsystems) equipped with night-vision goggles. The vitreous and pigment epithelium were removed, and the retina was isolated intact from the eyecup. After tension was released with four radial incisions, the dissected retina was attached to a membrane filter (AABG01300, Millipore Sigma) with a 3 mm hole in the center. The whole retina attached to the filter was mounted on a transparent imaging chamber (RC-26, Warner Instruments) with the photoreceptor layer side down. A slice anchor (SHD-26H, Warner Instruments) was placed on top of the retina. The chamber was mounted on a heated (30-32 °C) magnetic stage (PM-1, Warner Instruments) and continuously perfused with the same oxygenated Ames’ solution buffered with sodium bicarbonate throughout the experiments. The temperature was controlled by an automatic temperature controller (TC-344C, Warner Instruments).

### Visual stimuli

Visual stimuli were generated with custom Matlab (Mathworks) code on a Linux (Ubuntu, Canonical) machine with a GPU (AMD Radeon RX 580) using Psychophysics Toolbox extensions (Brainard, 1997) (Pelli, 1997). Stimuli were presented using a digital light processing (DLP) projector (PRO4500, Wintech), based on the LightCrafter 4500 (Texas Instruments), with an inserted ultraviolet (UV) LED (385 nm) and a frame rate of 60 Hz. The DLP was configured on a separate Windows (Microsoft) computer in a pixel-accurate mode (Pattern Sequence-Variable Exposure mode) to map each pixel to the native resolution (912 × 1140) of the digital micromirror device (DMD), bypassing its pre-programmed nonlinear video processing functions (scaling, gamma correction, and color coordinate adjustments). The pixels of the DLP were arranged in a diamond array configuration, so the coordinate scheme of the diamond pattern was mapped to a rectangular array of pixels (Marcireau et al., 2019), resulting in a 45-degree tilt angle in the output. The UV LED was used for visual stimulation, and the red LED output in spatially separated pixels was used to record the timing of visual stimulation. For each frame, the exposure time for red LED and UV LED was 0.8 ms and 15.5 ms, respectively. The size of a projected pixel of the DLP was 5.8 μm (0.17° FOV) on the retina, and the checker size was between 8 and 12 mm for spatiotemporal stimuli. Visual stimuli used in the experiments included full-field ON-OFF flashes (refresh rate: 0.167 Hz, duration: <1 min), randomly flickering checkerboards drawn from a Gaussian distribution (size: 1.5-2.2 mm, refresh rate: 6 Hz, duration: 30-40 min), and dynamic naturalistic movies from wildlife documentary clips (size: 2.2 mm, refresh rate: 30 Hz, duration: 30-40 min). The naturalistic stimuli were divided into 5-second blocks, and each block included transitions between multiple scenes. Each block was randomly rotated (90, 180 or 270 °) and/or flipped to avoid any directional motion biases within the original movie clips. The blocks were shuffled, and jittered with the statistics of eye movements.

### Two-photon Ca^2+^ imaging

#### Two-photon IR laser imaging

A custom-built two-photon laser scanning microscope was used for Ca^2+^ imaging. A femtosecond pulsed Ti:sapphire laser (Tsunami 3941-M1BB, Spectra-Physics) was tuned to 920-930 nm using a spectrometer (CCS175, Thorlabs). The beam passed through an electro-optic light modulator (350-105-02 and 302RM, Conoptics) and was raster-scanned with a resonant scanner system (RESSCAN and MDR-R, Sutter Instrument) using one resonant and one galvanometric mirror (CRS 8kHz and 6215, Cambridge Technology). The output was relayed to the back focal plane of the objective lens (HCX IRAPO L 25×, 0.95 NA W, 2.5 mm WD, Leica) through the scan lens (LSM54-1050, Thorlabs) and the tube lens (AC508-200-B, Thorlabs). The scan and tube lenses were in a telescope configuration, separated by the sum of their focal lengths. Laser excitation power was measured with an optical power meter (PM100USB, Thorlabs) and maintained at 10-20 mW at the back focal plane of the objective lens during the experiments. The optical components of the laser imaging path, including the objective lens, were mounted on a motorized platform (PK268M-E2.0B-C5, Oriental Motor) and moved in the x and y directions while the sample was stationary (Rosenegger et al., 2014). The objective lens also moved in the z direction. Translation was controlled by a motorized micromanipulator (MP-285A, Sutter Instrument). Emission light was directed to a long-pass dichroic mirror (FF775-Di01, Semrock), a long-pass filter (BLP01-488), and a convex lens (LA1708-A, Thorlabs). After another dichroic mirror (FF495-Di03, Semrock), a set of optical filters (FF01-496, FF01-790, FF01-538/84 and FF01-525/45, Semrock), and a convex lens (LA1805-A, Thorlabs), the emission light was collected with a photomultiplier tube (PMT) (H10769PA-40 and C8137-02, Hamamatsu Photonics). The output was amplified with a pulse amplifier (ZPUL-30P, Mini-Circuits) and connected to a timing discriminator (TD2000, Fast ComTec). The processed output was delayed with a digital delay generator (DB64, Stanford Research Systems) and adjusted to synchronize with the laser output pulses for photon counting. The output was sent to a channel of the digitizer adapter (NI-5734, National Instruments) of the data acquisition system (PXIe-6341 and PXIe-1073, National Instruments). The digitized signals were arranged and stored in an image stream (512 × 512 pixels) with a frame rate of 30 Hz in a recording computer using imaging software (ScanImage, Vidrio Technologies) written in Matlab. The size of the acquired image was equivalent to 0.6 mm^2^ on the retina at base zoom (1×), and the zoom factor was increased up to 2×.

#### Forward scattered light

The transmitted image of the tissue generated by the laser was captured to visualized non-fluorescent cells. The forward scattered laser light from the sample was directed through a convex lens to a photodiode (SM05PD1A and PBM42, Thorlabs) placed under the stage. The output was sent to one of the channels of the data acquisition system after a low-pass filter (DC-81 MHz, BLP-90+, Mini-Circuits).

#### IR LED imaging

A dichroic mirror (Di02-R980, Semrock) was attached to a motorized flipper (MFF101, Thorlabs) and positioned between the scan lens and the tube lens. The dichroic mirror was used to direct the transmitted light from the infrared (IR) LED (M940L3, Thorlabs) under the stage to a near infrared sensitive camera (GS3-U3-41C6NIR, Point Grey) for IR tissue imaging. The dichroic mirror was removed using the motorized flipper, and the IR LED was turned off (LEDD1B, Thorlabs) during two-photon imaging data acquisition.

#### Visual stimulation

The DLP projector was positioned below the stage and projected the stimulus through an objective lens (EC Plan-Neofluar 10×, 0.3 NA, 5.2 mm WD, Zeiss) from below. The red and UV channels were separated by a long-pass dichroic mirror (FF593-Di03, Semrock). The red channel was reduced by a neutral density filter (NE10B-A, Thorlabs), filtered by a long-pass filter (610ALP, Omega), collected by an achromatic lens (AC254-200-A, Thorlabs), and measured by a biased detector (DET36A2, Thorlabs). The output was amplified with a signal conditioner (Model 440, Brownlee Precision) and sent to one of the channels of the data acquisition system. The UV channel was further filtered by a set of optical filters (FF01-405/150 and FF01-468/SP, Thorlabs; ET376/30x, Chroma) and reduced by a neutral density filter (NDUV40B, Thorlabs). A dichroic mirror (FF580-FDi01, Semrock) was used to project visual stimuli through the objective lens onto the photoreceptor layer of the retina. The optical components for the visual stimulation were mounted on the same motorized platform as those for laser imaging, so that two-photon imaging and visual stimulation were mechanically aligned during translational movement of the imaging path in the x and y directions. The position of the visual stimulation pathway was further adjusted by an additional set of motorized actuators (CONEX-TRA25CC, Newport) prior to experiments.

### Data preprocessing

Fluorescence traces of each neuron were extracted from the collected data and transformed for further computational analysis. Regions of interest (ROIs) were detected from the recorded fluorescence signals by determining a threshold that minimizes the intra-class intensity variance of the binarized image and adjusting the number of connected pixels to be included using custom software written in Matlab. Further computational analysis was performed using custom Python code. The detected ROI traces were upsampled to 60 Hz and temporally aligned to the visual stimuli by using red LED signals measured by a photodiode (concurrently sampled with the PMT output). To reduce residual visual crosstalk noise directly from the DLP after spectral separation, a background subtraction was performed whereby the mean activity of two additional sets of adjacent background regions was subtracted from the ROI activity. These regions were in the direction of the line scan with the same number of pixels as the ROIs so that the background was sampled most closely in time with the ROIs and had similar variance to the ROIs. The average baseline of the extracted traces was calculated with a window size of 20 s, and traces were standardized with a window size of 30 s. The normalized traces were low pass filtered at 6 Hz and resampled to 30 Hz.

### Receptive fields

Spatiotemporal receptive fields were estimated by reverse correlation of the fluorescence response with spatiotemporal visual stimuli drawn from a Gaussian white noise distribution. Visual stimuli were resampled in the time domain to 30 Hz, and the fluorescence response was converted to the same sampling rate. The reverse correlation of fluorescence responses *F*′(*t*) with stimulus *s*(*t*) was calculated as

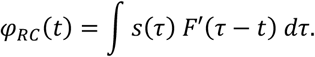

Spatial and temporal receptive fields for display were obtained by singular value decomposition of the spatiotemporal receptive field matrix (*φ*) by *φ* = *U* Σ *V*^*T*^. The first column vectors of *U* and *V* were used to approximate the temporal and spatial receptive fields (rank-1 approximation). The spatial dimension was reduced to one before decomposition and recovered from the first column of the decomposed spatial receptive field.

#### Convolutional Neural Network (CNN) models

We trained Convolutional Neural Network (CNN) models to predict the fluorescence responses of neurons to naturalistic movies, similarly as previously described (McIntosh et al., 2016) (Maheswaranathan et al., 2023). Four datasets from different experients were used for model training. Among hundreds of ROIs (188, 137, 227, and 283 for each dataset), accepted ROIs with a sufficiently high correlation between repeats of the same stimulus were selected for training datasets. The datasets consisted of 75, 75, 60, and 75 ROI responses from fluorescence recordings with dynamic natural scenes stimulus. A CNN model was trained for each dataset. Model parameters were optimized to minimize a loss function corresponding to the mean squared error (MSE) between the data and model predictions,

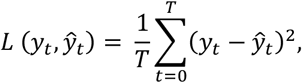

where *y*_*t*_ is the measured response and *ŷ*_*t*_ is the predicted model output at time *t. T* is the batch size and equivalent to 2000 samples. All model parameters were first optimized to simultaneously fit 30 selected ROI responses with a high reliability across trials, by minimizing the MSE between the selected ROI responses and corresponding model outputs. After initial model training, the model parameters were saved and used for fine-tuning with the other 60-75 ROI responses, including the initial 30 ROIs. During the fine-tuning stage, the model parameters of the first layer were set to the learned parameters from the pretrained model, and only parameters of the second layer were trained.

The architecture of the CNN models consisted of one convolutional layer (Layer 1, 17 x 17 pixel), with 6 channels, followed by a fully connected layer (Layer 2, 16 x 16 pixel). Each convolutional and fully connected layers consisted of a linear filter, a batch normalization layer, and a nonlinear softplus function. After the last softplus function, an affine layer was added for scaling. The input of the model was a tensor with size of *N*_*train*_ x 40 x 32 x 32 (# of samples x time x space x space), and the output of the model was a tensor with size of *N*_*train*_ x *N*_*ROI*_ (# of samples x # of ROIs), where *N*_*train*_ is the number of training samples (53940-71930) and *N*_*ROI*_ is the number of ROIs for model training (60-75). Model performance was evaluated by comparing validation/test dataset and corresponding model output, and the number of samples for validation was between 1790-2990. The validation dataset was an average response from 5-6 repeats of an identical stimulus, which is different and stored separately from the training stimulus. The models were trained for up to 200 epochs, and the training was stopped when validation accuracy decreased to avoid overfitting of the parameters to training dataset. The learning rate (0.001) and L2 regularization hyperparameter (0.01) were chosen based on the model’s performance.

Model optimization was performed using the Adam optimization algorithm, an extension to stochastic gradient descent with adaptive properties. Models were trained using PyTorch on Macbook Air (M1, Apple).

### Interneuron contributions

To compute how much each interneuron contributes to the output responses of the neurons, we applied Integrated Gradients (IG) (Sundararajan et al., 2017). To decompose the model output into attributes for each interneuron activity (Dhamdhere et al., n.d.) (Leino et al., 2018) in response to the input movie, we computed Interneuron Contributions (INCs) by applying the chain rule as previously described (Tanaka et al., 2019) (Maheswaranathan et al., 2023).

We applied the chain rule to decompose the responses of amacrine cells into INCs for each of first layer model units. Formally, the trained CNN model is a function *r*(*t*) = *F*[*s*(*t*)], where *r*(*t*) is the model output, *s*(*t*) is the stimulus input, and *F* is the nonlinear function. By using path integration, *s*(*t*, α) = α · *s*(*t*), where the path takes a straight line from α = 0 (gray background) to α = 1 (actual stimulus presented), the model output can be represented as,

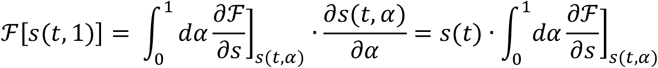

We considered contributions of the rectified output of the Layer 1 model units, whose responses to the stimulus are 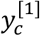, where [1] refers to an index of the layer and *c* refers to channel or model cell type. We can define the INC of the *c*-th channel 𝒴_*c*_ *as*,

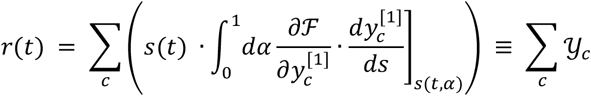

When the INCs, 𝒴_*c*_, were spatially averaged, the resulting vector was with six elements at each time point, representing the contribution of the rectified output of each Layer 1 channel to the final output.

### Clustering

#### k-means clustering

The amacrine cells were grouped by their averaged flash responses by using standard *k*-means clustering algorithms. The number of clusters, *k*, was chosen based on the change of variance explained by varying the number of clusters.

#### Hierarchical clustering

We clustered amacrine cells by their INCs from rectified output of Layer 1 under full-field flash or natural scenes by using hierarchical clustering. At each iteration, two clusters were combined by using the Ward variance minimization algorithm, and the distance of the newly formed cluster with the remaining clusters was computed. The spatially averaged INCs matrix was normalized and sampled every 5-20 frames, yielding a vector size of 6 x *N*_*t*_ (# of channels x # of samples) per each ROI. The vectors were used to compute the Euclidean distance between clusters. The clustered amacrine cells were visualized in the low-dimensional space by the t-distributed Stochastic Neighbor Embedding (t-SNE).

After clustering of amacrine cells, the INCs within the same cluster during flash stimulus were averaged. The vectors for each cluster were appended and 42 (= 6 x 7) elements were computed for each time point (# of channels x # of clusters). This element represents the state of the entire network at different time. We performed the same clustering method to this matrix to identify stimulus-dependent states of the network at each point in time.

## Acknowledgments

The authors wish to acknowledge members of the Baccus laboratory for helpful discussions and technical assistance. We would like to thank Sarah E. Ross for providing Ptf1a-cre mice, and Alexandre Marcireau and Ben Smith for technical assistance. This work was supported by grants from the NEI (R01EY022933, R01EY025087, P30-EY026877) (SAB).

## Author Contributions

All authors participated in the overall design of the study and wrote the manuscript. D.L. and J.K. performed experiments and analyzed data.

## Supplemental Information

**Supplemental Figure 1.**
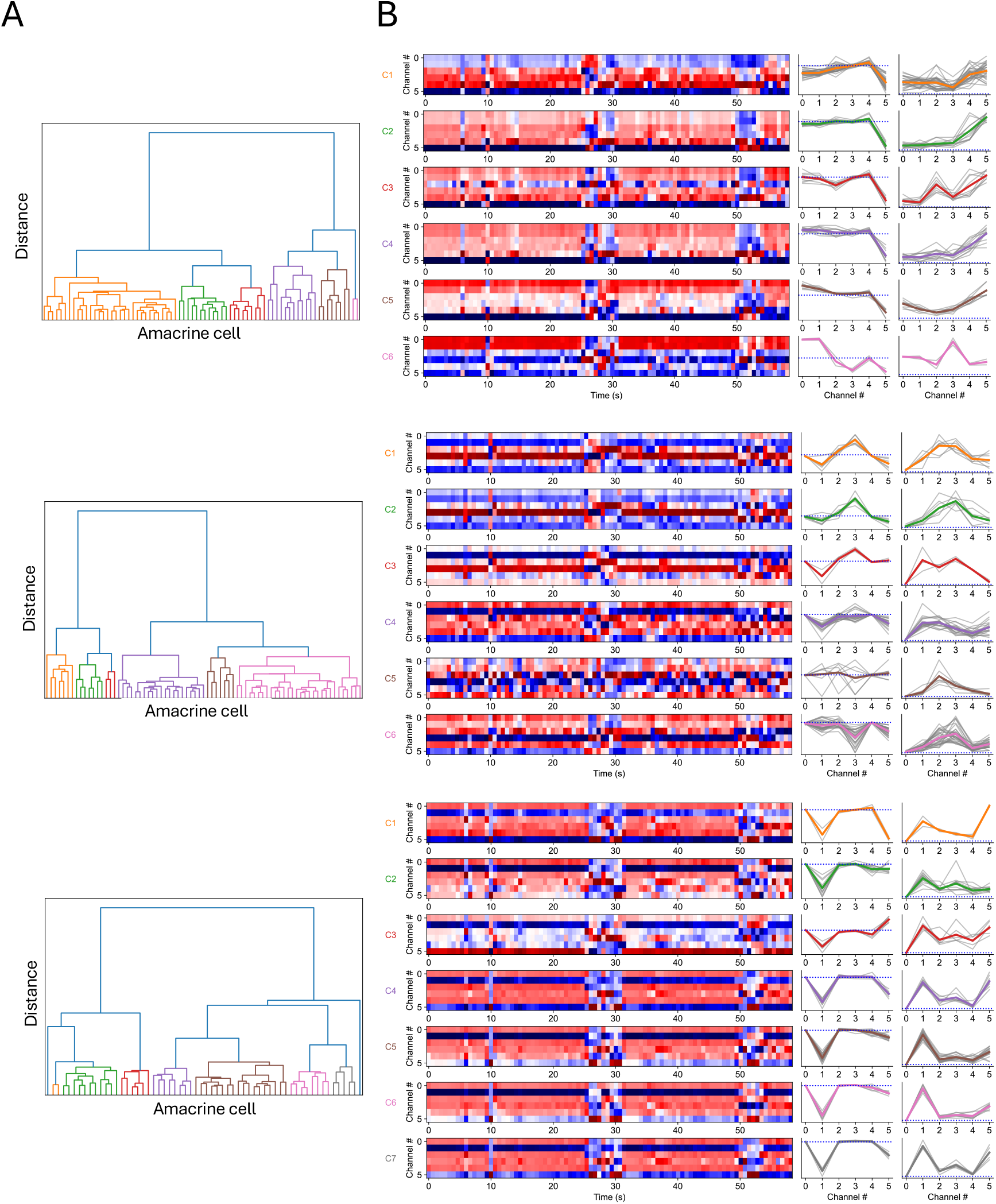
Examples of interneuron contributions for different preparations. Shown are three additional preparations analyzed as in Fig. 4. A. Hierarchical clustering diagram of Layer 1 INCs that caused amacrine cell responses to natural scenes as in (Fig. 4C). B. INCs for each amacrine cell cluster in response to a 60 s segment of natural scenes as in (Fig 4D). Left. Each row within each amacrine clusters response is a different Layer 1 cell type (Channels 0 – 5). Red indicates excitation, blue indicates inhibition. Middle. Average INC for each Layer 1 cell type for each cluster. Right. Standard deviation of INC for each cluster, showing the magnitude of the contribution of each Layer 1 cell to each amacrine cluster.

## References

Baden, T., Berens, P., Franke, K., Rosón, M.R., Bethge, M., Euler, T., 2016. The functional diversity of retinal ganglion cells in the mouse. Nature 529, 345–350. doi:10.1038/nature16468

Brainard, D.H., 1997. The psychophysics toolbox. Spat Vis 10.

Dhamdhere, K., Sundararajan, M., 1805.12233, Q.Y.A.P.A., 2018, How important is a neuron?, arXiv.

Gollisch, T., Meister, M., 2010. Eye smarter than scientists believed: neural computations in circuits of the retina. Neuron 65, 150–164. doi:10.1016/j.neuron.2009.12.009

Helmstaedter, M., Briggman, K.L., Turaga, S.C., Jain, V., Seung, H.S., Denk, W., 2013. Connectomic reconstruction of the inner plexiform layer in the mouse retina. Nature 500, 168–174. doi:10.1038/nature12346

Kolb, H., Dekorver, L., 1991. Midget ganglion cells of the parafovea of the human retina: a study by electron microscopy and serial section reconstructions. J Comp Neurol 303, 617–636. doi:10.1002/cne.903030408

Leino, K., Sen, S., international, A.D.2.I., 2018, Influence-directed explanations for deep convolutional networks. https://ieeexplore.ieee.org

Maheswaranathan, N., McIntosh, L.T., Tanaka, H., Grant, S., Kastner, D.B., Melander, J.B., Nayebi, A., Brezovec, L.E., Wang, J.H., Ganguli, S., Baccus, S.A., 2023. Interpreting the retinal neural code for natural scenes: From computations to neurons. Neuron. doi:10.1016/j.neuron.2023.06.007

Marcireau, A., Ieng, S.-H., Benosman, R., 2019. Sepia, Tarsier, and Chameleon: A Modular C++ Framework for Event-Based Computer Vision. Front Neurosci 13, 1338. doi:10.3389/fnins.2019.01338

Masland, R.H., 2012. The tasks of amacrine cells. Vis. Neurosci. 29, 3–9.

McIntosh, L.T., Maheswaranathan, N., Nayebi, A., Ganguli, S., Baccus, S.A., 2016. Deep Learning Models of the Retinal Response to Natural Scenes. Adv Neural Inf Process Syst 29, 1369–1377.

Pelli, D.G., 1997. The VideoToolbox software for visual psychophysics: transforming numbers into movies. Spat Vis 10, 437–442.

Segev, R., Puchalla, J., Berry, M.J., 2006. Functional organization of ganglion cells in the salamander retina. J. Neurophysiol. 95, 2277–2292. doi:10.1152/jn.00928.2005

Sundararajan, M., Taly, A., Yan, Q., 2017. Axiomatic attribution for deep networks, in:. Presented at the Proceedings of the th International Conference on Machine Learning, pp. 3319–3328.

Tanaka, H., Nayebi, A., Maheswaranathan, N., McIntosh, L., Baccus, S., Ganguli, S., 2019. From deep learning to mechanistic understanding in neuroscience: the structure of retinal prediction, in: Wallach, H., Larochelle, H., Beygelzimer, A., Buc, F.D.A.E., Fox, E., Garnett, R. (Eds.). Presented at the Advances in Neural Information Processing Systems, Curran Associates, Inc.

Vaney, D.I., Young, H.M., Gynther, I.C., 1991. The rod circuit in the rabbit retina. Vis. Neurosci. 7, 141–154.

Yan, W., Laboulaye, M.A., Tran, N.M., Whitney, I.E., Benhar, I., Sanes, J.R., 2020. Mouse Retinal Cell Atlas: Molecular Identification of over Sixty Amacrine Cell Types. J. Neurosci. 40, 5177–5195. doi:10.1523/JNEUROSCI.0471-20.2020

